# The RNase and RNA binding activities of selected RNase R truncations and mutations plus a detailed step by step protocol to purify recombinant RNase R

**DOI:** 10.64898/2026.04.15.718802

**Authors:** Wataru Horikawa, Daniel L. Kiss

## Abstract

RNase R is a processive 3’ to 5’ exoribonuclease that degrades a broad array of linear RNA species while preserving RNA lariats and circular RNAs (circRNA). In recent years, this enzyme has become pivotal for the field of circRNA research, serving as a key step for circRNA enrichment, purification, and identification. Despite this growing importance, the effects of mutations and truncations in RNase R have been incompletely studied. We make several point mutations and assay their effects on the ability of RNase R to bind and/or degrade RNA substrates. Our data show that selected active site mutations have varying effects on RNA binding and degradation.

Furthermore, the increasing interest in circRNA-based RNA therapeutic platforms highlights an urgent need for RNase R in RNA molecular biology labs. However, the substantial cost of commercial RNase R remains a bottleneck, particularly for large-scale studies or the development of circRNA-based technologies. In this protocol, we offer a solution to that problem, namely a more accessible and cost-effective means of purifying high-quality and low-cost RNase R. We provide a highly detailed yet simplified, high-yield protocol that produces recombinant RNase R from *Escherichia coli*. The method uses a single-step Ni-NTA affinity chromatography procedure without proteolytic tag removal and is optimized for entry-level FPLC systems such as the ÄKTA Start, ensuring that high-purity enzyme production does not require specialized, high-end instrumentation. A second key feature is the establishment of an optimized reaction framework, including specific buffer compositions and defined enzyme-to-substrate ratios for the purified RNase R. The protocol achieves functional equivalence to premium commercial RNase R, ensuring complete linear RNA digestion without compromising the integrity of circRNA. The combination of a simplified purification workflow and a robust reaction protocol provides an accessible, cost-effective, and reliable solution for any molecular biology laboratory requiring high volumes of RNase R.

**Key Features:**

- RNase R mutations can block RNase activity, RNA binding or both
- This protocol purifies ∼40 mg of active RNase R per liter of E. coli culture
- The protocol avoids medium and high end FPLC systems
- RNase R expression constructs (WT and mutants) will be available on Addgene
- The protocol includes an optimized reaction buffer to pair with this RNase R
- Optional endotoxin removal step is also included

## Introduction

Ribonucleases such as RNase A, RNase T1, and RNase R have been key reagents in RNA biology research for decades. Notably, RNase R, a highly processive 3’-to-5’ exoribonuclease capable of unwinding and degrading highly complex RNA sequences including secondary structures that can arrest other RNases, has become even more important [1-3]. RNase R only requires that the ‘target’ RNA have a free 3’ terminus for it to be recognized and degraded. Meaning that any (and all) manner of linear RNA species can be eliminated with RNase R digestion, but intron lariats and circular RNAs (circRNAs) cannot. This specificity for linear RNAs has made RNase R a key tool in circRNA research [4-6].

Despite its utility, commercial RNase R can be prohibitively expensive and can limit the ability of some teams doing select types of RNA-focused research (like some teams working with circRNAs). While several purification methods have been published, they are often complex and costly. This protocol is a streamlined method to purify recombinant His-tagged RNase R and generally yields ∼40 mg of purified protein per liter of *E. coli* culture. We have also optimized the reaction buffer and RNA digestion conditions to make RNase R that is functionally indistinguishable from commercially sourced RNase R.

As mentioned above, RNase R has become a critical reagent for studying circRNAs. Its ability to recognize and degrade only linear RNAs is one key reason for this. We hypothesized that we could harness that trait to capture linear RNAs in a degradation independent manner. Specifically, we reasoned that a catalytically inactive RNase R mutant could bind, but not degrade linear RNAs, while leaving circRNAs untouched. We constructed and tested multiple versions of RNase R to test this hypothesis.

## Results and discussion

We used traditional cloning methods to create bacterial expression plasmids encoding different wild type (WT) or mutant RNase R proteins. **Table 1** shows the plasmid constructs used in this study.

**Table 1:** Bacterial expression plasmid constructs made for this study. The Addgene numbers, the identity of the expressed protein, and the sizes of the insert, plasmid, and plasmid backbone minus the insert are listed in base pairs (bp) for each construct.

| Addgene number | Insert | insert length (bp) | plasmid length (bp) | backbone only (bp) |
| --- | --- | --- | --- | --- |
| 254212 | His6-V5-RNaseR-87-725-D280N | 2019 | 7677 | 5658 |
| 254213 | His6-V5-RNaseR-87-725 | 2019 | 7677 | 5658 |
| 254214 | His6-V5-RNaseR-1-725 | 2541 | 8199 | 5658 |
| 254215 | His6-V5-RNaseR-1-725-D280N | 2541 | 8199 | 5658 |
| 254216 | His6-V5-RNaseR-87-725-D278N | 2019 | 7677 | 5658 |
| 254217 | His6-V5-RNaseR-87-725-D280A | 2019 | 7677 | 5658 |
| 254218 | His6-V5-RNaseR-87-725-D278A | 2019 | 7677 | 5658 |

### Purification of WT and mutant RNase R proteins

The plasmids were transformed into *E. coli* and the proteins were purified as described in detail below in the extended Materials and Methods section. The integrity of purified WT and mutant RNase R proteins were assessed using Stain-free SDS PAGE gels (**Figure 1**) where the asterisks mark the expressed protein. The His6-V5-RNaseR-87-725 is the most efficient version of the RNase R protein we purified, and the protocol is optimized to purify this protein. Notably, this version of RNase R has truncations at both the N-terminus (Δ1-86) and the C-terminus (Δ726-813). We were able to express and purify an N-terminally intact (His6-V5-RNaseR-1-725) version of RNase R; however, we found it prone to precipitation which made it challenging to use as a reagent. In our hands, it always precipitated once we added it into our reaction buffer. With enough optimization, you may find conditions where it can be used, but since the 87-725 double truncation has robust activity and is much easier to purify, we strongly recommend using that form of RNase R. Notably, we were never able to purify RNase R D280N and the RNase defective null mutants without some level of nucleic acid contamination. Regardless of many tested conditions, as judged by A260/A280 ratios, our preparations of the 1-725-D280N, 87-725-D280N, 87-725-D280A, 87-725-D278N, and 87-725-D278A mutants were always contaminated with nucleic acids ∼likely bacterial RNAs∼ bound during protein expression [7]. This included attempts to purify the mutant RNase R proteins in the presence of urea to denature the protein and RNA (data not shown).

**Figure 1:**
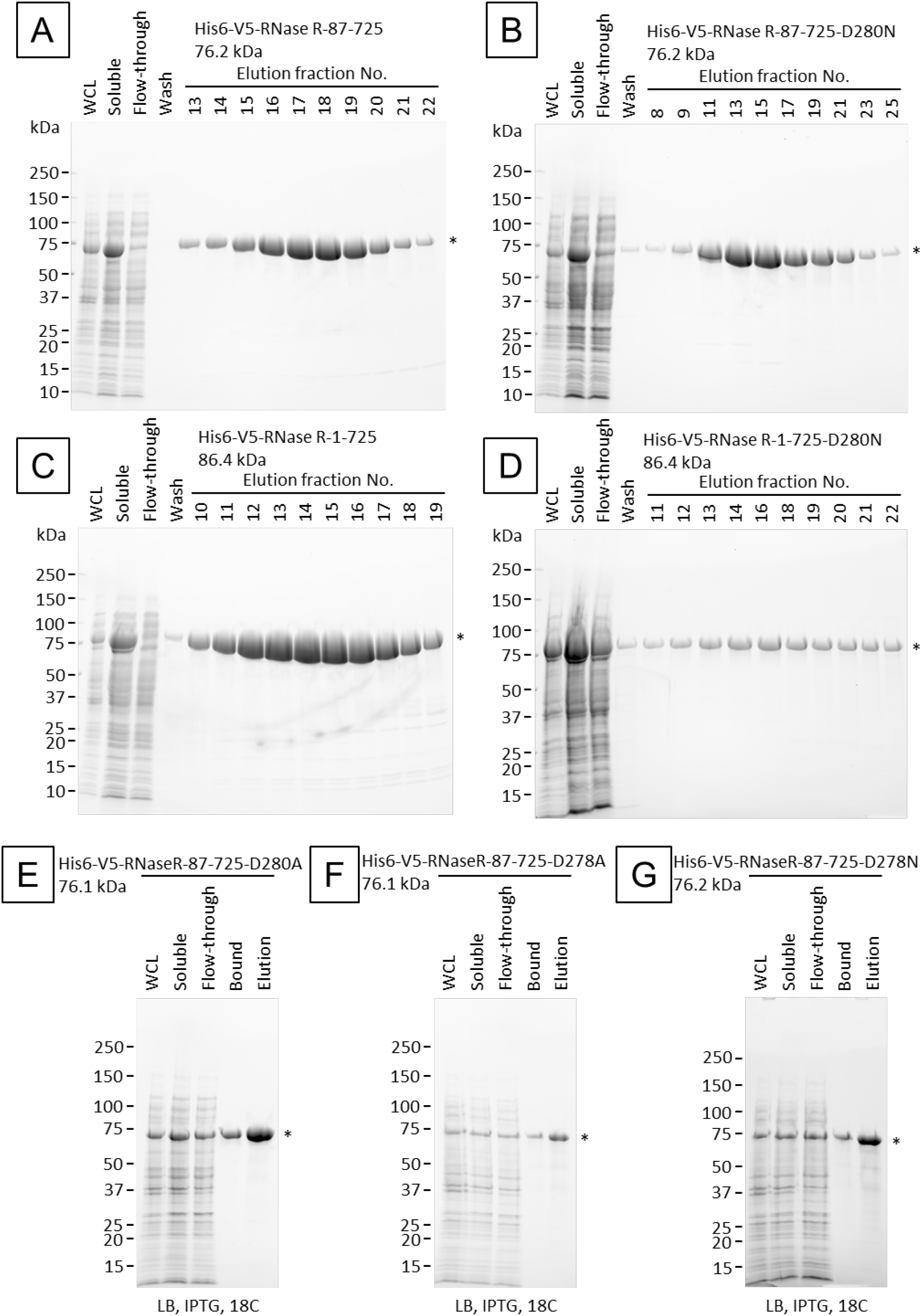
Purification of wild type and mutant RNase R proteins. Fractions were collected at different parts of the protein purification protocol. These include whole *E. coli* cell lysates (WCL), flowthrough samples, bead-retained (bound), and eluted fractions. Aliquots of each fraction were denatured in SDS loading dye and loaded onto stain-free Any kD SDS PAGE gels. Proteins were visualized with a Chemi doc. Selected elution fractions of purified **(A)** RNase R-87-725, **(B)** RNase R-87-725-D280N, **(C)** RNase R-1-725, **(D)** RNase R-1-725-D280N, or final purified and dialyzed fractions for **(E)** RNase R-87-725-D280A, **(F)** RNase R-87-725-D278A, **(G)** RNase R-87-725-D278N. The gels shown are representative examples of between one and six independent replicates.

### Exoribonuclease activities of WT and mutant RNase R proteins

The purified proteins were tested for their ability to degrade different RNA substrates. The WT RNaseR-87-725 efficiently degrades a 25-mer RNA oligonucleotide when the RNA and protein were at a 1:1 ratio (**Figure 2A, 2B**). Furthermore, we see that either of two mutations, D280N or D280A, fully block the RNase activity or the protein; however, two other mutations D278A and D278N only partially abrogate the protein’s RNase activity (**Figure 2A, 2B**). Although predicted to be an inactivating mutation, unlike with D280N (which was a true null), we still observed some RNase activity with both D278 mutations. D278A was ∼slightly∼ more active than D278N and we estimate they both retained ∼35-50% of WT activity (**Figure 2A, 2B**). We also validated that our WT RNaseR-87-725 could specifically degrade long mRNAs while leaving circular RNAs (circRNAs) intact (**Figure 2C, 2D**). Please note that RNase R purified with this protocol requires a special reaction buffer, and is not active in at least three buffers provided with RNase R from three different commercial sources (data not shown). Lastly, we compared the activity and specificity of our RNase R against commercially-supplied RNase R. As shown in **Figure 2D**, we show similar activity and specificity for linear RNAs as commercially sourced RNase R.

**Figure 2:**
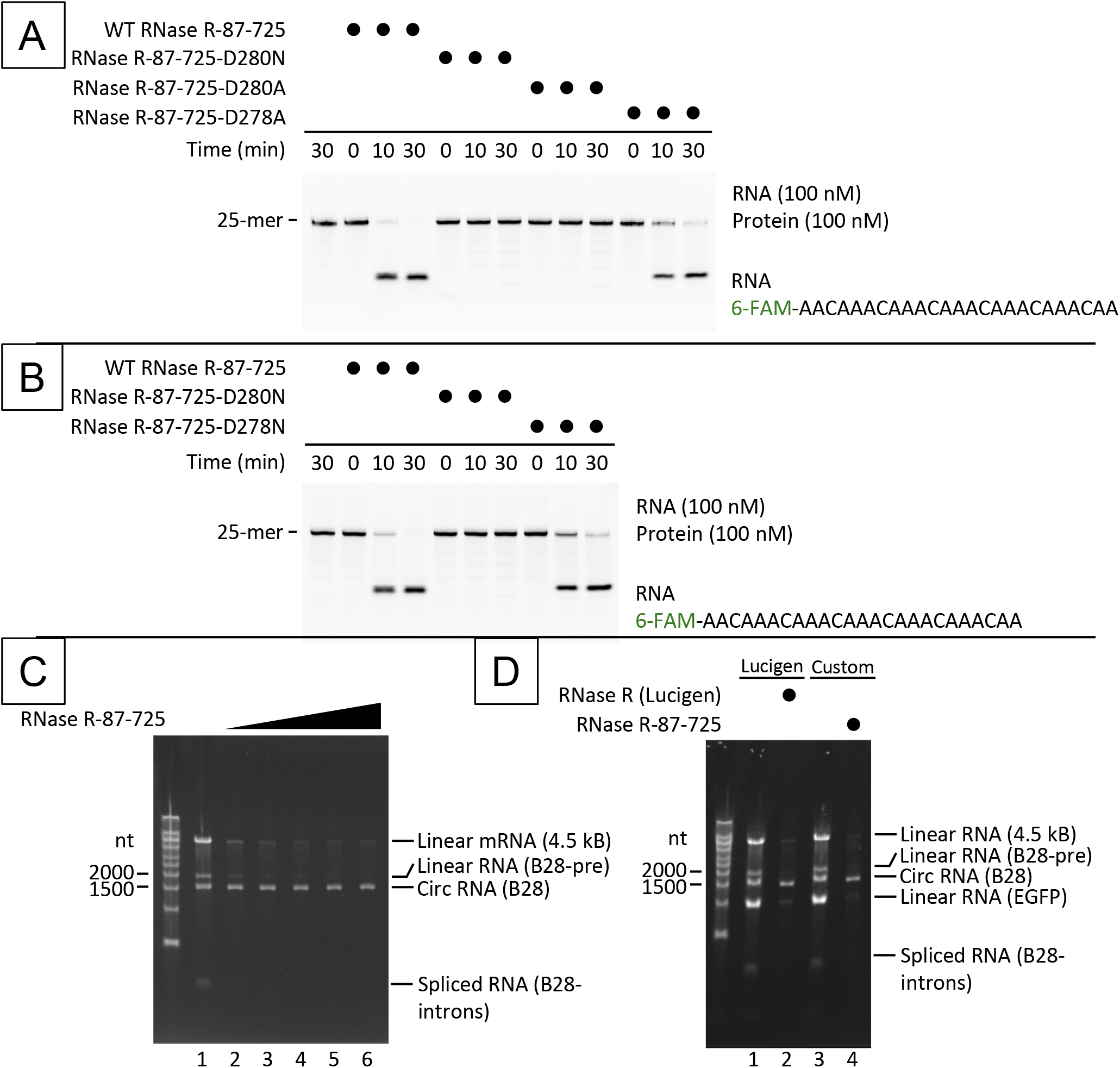
RNase activity of wild type and mutant RNase R proteins. The RNase activities of purified **(A)** WT RNase R-87-725 (5 replicates), RNase R-87-725-D280N (2 replicates), RNase R-87-725-D280A (1 replicate), and RNase R-87-725-D278A (1 replicate) and **(B)** RNase R-87-725-D280N (2 replicates) plus RNase R-87-725-D278N (1 replicate) on a FAM-labeled 25-mer RNA oligo. **(C)** RNase activity of WT RNase R-87-725 on a mixture of linear mRNAs (B28-pre, 4.5 kB, spliced introns) and circular RNA (B28), **(D)** RNase activity of WT RNase R-87-725 on a mixture of linear mRNAs (B28-pre, 4.5 kB, EGFP mRNA, and spliced introns) and circular RNA (B28) compared to a commercially-sourced RNase R (Lucigen). The buffer systems are: Lucigen: 20 mM Tris-HCl (pH 7.5), 100 mM NaCl, 0.25 mM MgCl_2_, 1 mM DTT and custom: 20 mM Tris-HCl (pH 7.5), 100 mM KCl, 0.25 mM MgCl_2_. The data shown in panels **(C)** and **(D)** are representative examples of three or more independent experiments.

### RNA binding abilities of select RNase R mutants

The purified proteins were evaluated for their ability to bind a small RNA substrate via electromobility shift assays (EMSA). As the WT enzymes were shown to almost completely degrade these small RNA substrates in ∼10 minutes, we did not attempt to use them in this set of experiments. Three of the point mutations in the RNaseR-87-725 construct showed no RNA binding ability (**Figure 3A-C**). Although the RNase R-87-725-D280A was completely null in its ability to degrade the labeled RNA, it did show some ability to bind the RNA oligo when the protein was in great excess (**Figure 3D**). We also tested the RNA binding ability of the D280N mutant when the enzyme had an intact N-terminus and saw that it was barely able to bind RNA (**Figure 3E**) at highly elevated protein:RNA ratios (<5% of RNA bound with a ratio of 10:1 protein:RNA). This did increase to ∼15% when the reactions were briefly UV crosslinked (**Figure 3F**) suggesting transient or weak binding. However, even weak binding was an unexpected result since our N-terminally-truncated D280N mutant showed absolutely no RNA binding in this assay. These data could suggest that the N-terminus of RNase R has its own RNA binding function.

**Figure 3:**
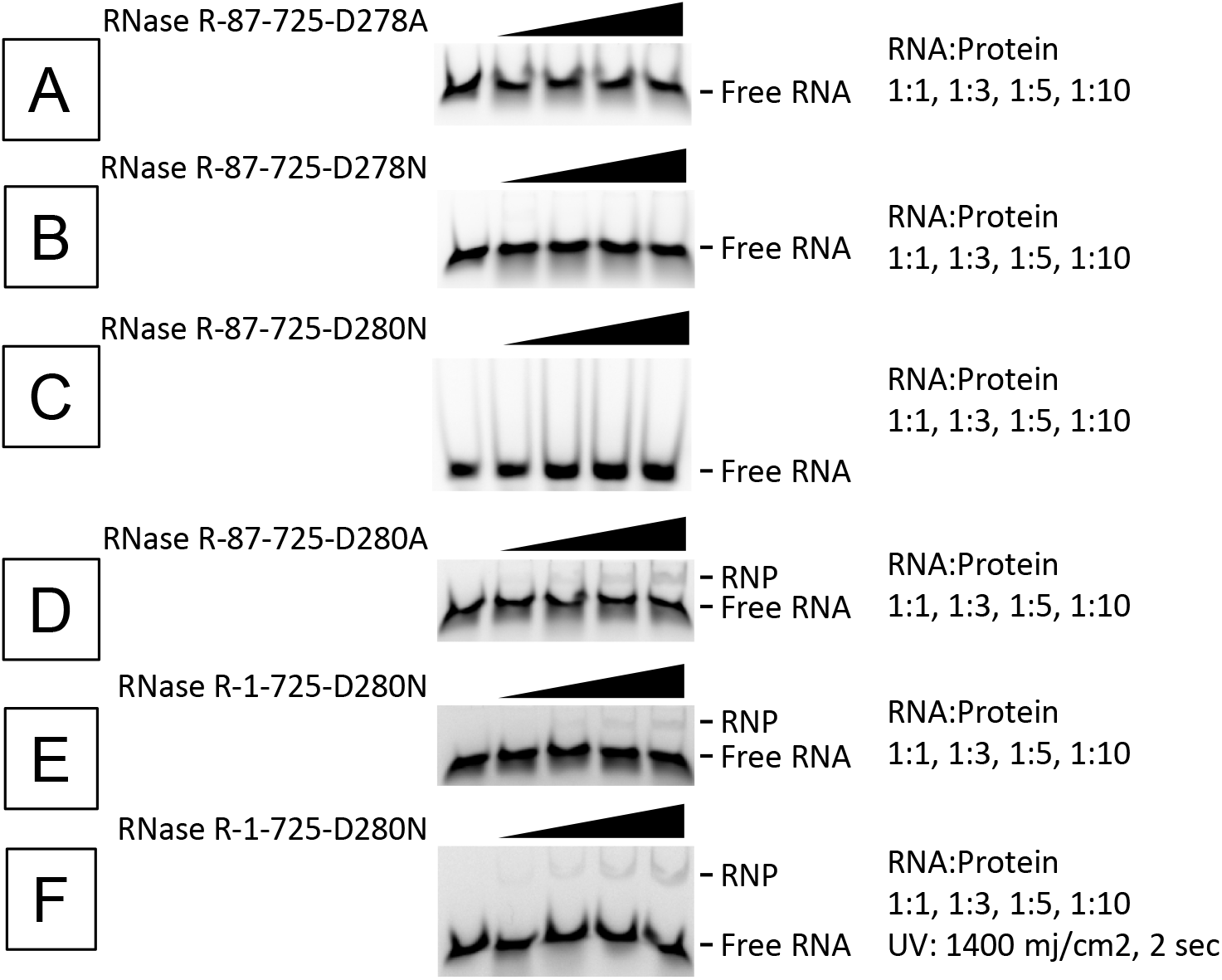
RNA binding ability of mutant RNase R proteins. Electromobility shift assays (EMSA) showing the RNA binding abilities of purified RNase R mutant proteins. A 5’FAM-labeled 25-mer RNA oligonucleotide was mixed with the indicated RNase R mutant using the indicated RNA:protein ratios. The complexes were separated using native polyacrylamide gels and visualized using the FAM label and a ChemiDoc. The population of shifted RNA bands are labeled as RNP, while unbound RNA is marked as free RNA. The mutants used are **(A)** RNase R-87-725-D278A (1 replicate), **(B)** RNase R-87-725-D278N (1 replicate), **(C)** RNase R-87-725-D280N (4 replicates), **(D)** RNase R-87-725-D280A (2 replicates), and RNase R-1-725-D280N either without **(E)** or with **(F)** UV crosslinking (1 replicate each).

The data shown above strongly disfavor our initial hypothesis that inactivated RNase R can be used to efficiently capture linear RNAs. While we did observe some RNA binding with selected RNase R mutations, this only occurred at impractical ratios of RNase R protein:RNA target. However, our inability to purify nucleic acid-free RNase R mutants is a key limitation of this work. Simply speaking, our true RNase R null mutations (like D280N) behave like stereotypical ‘substrate traps.’ Meaning that they may bind their target substrate so tightly that they are unable to release them. One possible interpretation of our data is that our in vitro assays were doomed to fail since our nuclease dead mutants already had RNAs (from *E. coli*) bound in their active sites which prevented the proteins from holding our test RNAs.

### COMMENTARY

Over the course of performing the research detailed above, we realized that the RNase R purification protocol itself would be a very useful tool for the broader RNA biology research community. As such, we endeavored to produce a highly detailed, easy to follow method so other teams could use our plasmids to purify their own RNase R. This protocol delineates the procedures to express, purify, and ensure the solubility of active RNase R. The typical yield of RNase R produced by this protocol is roughly 40 mg of protein per liter of LB medium. However, the protein yield may vary based on several factors including the quality of the transformed cells, the type of medium employed (we have found that lower quality/inexpensive LB from less reputable suppliers reduces yields), the size of cell pellets at the start points and the condition of the cells. Therefore, it is advisable to use freshly transformed cells. In addition, we recommend expediting the protein purification process to avoid protein degradation. Although we recommend moving quickly through the protocol once cell lysis has begun, the RNase R purified using this protocol has shown to be quite stable in storage with no detectable loss of activity despite 14 months of storage at -20°C (data not shown).

## Materials and Methods

### Basic Protocol 1

#### Purification of RNase R from *Escherichia coli*

This protocol details the purification of RNase R with N-terminal 6x His-tag (hexa-histidine tag) using BL21 Star (DE3) cells. We use a nickel-affinity chromatography column and an ÄKTA start FPLC system to isolate the recombinant protein. This purification protocol takes about 3 days and yields around 40 mg of RNase R per liter of LB medium.

#### Materials

##### Please note

We list our preferred reagents and catalog numbers here. However, (with very few exceptions) we are certain that equivalent items can be substituted according to lab preferences and availability without loss of protein yield or activity. With that said, we do note certain items where reagent quality (the LB medium for example) is directly linked to protein yields. Please keep this in mind when gathering materials and performing this protocol.

- Plasmid DNA encoding *E. coli* Rnase R (**Please note:** To facilitate distribution, the plasmids mentioned in this protocol will be available through Addgene under accession numbers #254212 – #255218. This protocol is optimized for #254213, a fully active N- and C-terminally truncated form of RNase R that remains stable in solution.)
- One Shot BL21 Star (DE3) Chemically Competent Cells (Invitrogen, REF: 44-0049)
- LB medium (Lennox) (see recipe)
- LB agar (1.2%) plate with appropriate antibiotics (BD, REF: 240230; Apex Bioresearch, Cat#: 20-273)
- Ampicillin, 100 mg/ml (Sigma-Aldrich, Cat#: A5354-10ML)
- SOC Outgrowth Medium (New England Biolabs, Cat#: B9020S)
- IPTG stock solution, 1 M (see recipe)
- Antifoam 204 (Sigma-Aldrich, Cat#: A6426)
- Lysis buffer (see recipe)
- Optional: Endotoxin removal wash buffer (see recipe)
- Wash buffer (see recipe)
- Elution buffer (see recipe)
- Dialysis buffer (see recipe)
- HisTrap HP 5 mL (Cytiva, Cat#: 17524802)
- 2x Laemmli Sample Buffer (Bio-Rad, Cat#: 1610737)
- Precision Plus Protein Unstained Standards (Bio-Rad, Cat#: 1610363)
- Mini-PROTEAN TGX Stain-Free Precast Gels, 4-20%, 15-well comb (Bio-Rad, Cat#: 4568096)
- (alternate) appropriate mixtures of 29:1 acrylamide, ammonium persulfate, and TEMED for making SDS PAGE gels
- Spectra/Por 6 Dialysis Membrane Pre-wetted RC Tubing, MWCO: 3.5 kDa (Spectrum, REF: 132592)
- Spectra/Por Closures Standard Type, 55 mm (Spectrum, REF: 132737)
- Spectra/Por Closures Weighted, 55 mm (Spectrum, REF: 132745)
- Slide-A-Lyzer Buoys (Thermo Scientific, REF: 66432)
- MCE Membrane Filter, 0.22 μm Pore Size (Millipore, REF: GSWP04700)
- RNA Flashgel (Lonza)
- (alternate) appropriate mixtures of 19:1 acrylamide, ammonium persulfate, TEMED, and urea for making urea PAGE gels
- 2-, 10-, 20-, 200-, and 1000-μl pipets
- 10-, 200-, and 1000-μl pipet tips
- 250 mL Erlenmeyer flask (VWR, Cat#: 10536-914) or equivalent
- Petri dish, 90 mm diameter (Genesee Scientific, Cat#: 32-107G) or equivalent
- 15 ml centrifuge tube (Olympus Conical, Cat#: 28-103) or equivalent
- 50 ml centrifuge tube (Olympus Conical, Cat#: 28-108) or equivalent
- 2.5 L Full-Baffle TUNAIR Polypropylene Erlenmeyer Flask (IBI SCIENTUIFIC) or equivalent
- Dri-gauz filter liner for TUNAIR shake flask (IBI SCIENTUIFIC)
- 1 L (1000 mL) Polypropylene Bottle Assembly (Beckman Coulter, Product No: C31597) or equivalent
- 50 mL, Polypropylene Bottle with Cap Assembly (Beckman Coulter, Product No: 361694) or equivalent
- Nalgene Reusable Bottle Top Filters (Thermo Scientific) or equivalent
- Disposable Graduated Transfer Pipets (VWR, Cat#: 414004-017) or equivalent
- Silicone spatula
- 250 mL, 500 mL, and 1 L graduated cylinders
- 250 mL, 500 mL, and 1 L beakers

#### Required Equipment

- Autoclave
- -80°C freezer
- -20°C freezer
- 4°C cold room, chromatography refrigerator, or refrigerator with power for a magnetic stir plate
- 37°C incubator for bacterial plates
- Magnetic stir plate (Benchmark Scientific) or equivalent
- Balance (Benchmark Scientific) or equivalent
- Innova S44i Shaker (Eppendorf) or equivalent
- Eppendorf® Lab Shaker Clamps (Eppendorf, Cat #: M1190-9005) or equivalent
- Eppendorf® 5424 refrigerated microcentrifuge or equivalent
- Allegra X-30R Benchtop Centrifuge (Beckman) or equivalent
- Avanti JXN-26 High speed centrifuge (Beckman Coulter) or equivalent
- J-LITE JLA-8.1000 Fixed-Angle Aluminum Rotor (Beckman Coulter, Product No: 363688) or equivalent
- JA-25.50 Fixed-Angle Rotor (Beckman Coulter, Product No: 363055) or equivalent
- Pulse 150 UltraSonic Homogenizer (Benchmark Scientific) or equivalent
- 6mm horn for Pulse 150 and Pulse 650 (Benchmark Scientific) or equivalent
- ÄKTA start FPLC system and Frac30 fraction collector (Cytiva) or equivalent
- UNICORN start v1.3 software (Cytiva) or equivalent
- MultiTherm Shaker with heating and cooling (Benchmark Scientific) or equivalent
- ChemiDoc Imaging System (Bio-Rad) or equivalent
- Mini-PROTEAN Tetra Vertical Electrophoresis Cell (Bio-Rad) or equivalent
- PowerPac Basic Power Supply (Bio-Rad) or equivalent
- Spectrophotometer (DeNovix, Cat#: DS-11 FX+) or equivalent
- Vortex mixer (Benchmark Scientific, Cat#: BV1000) or equivalent
- Flashgel dock (Lonza) or equivalent
- Optional: Masterflex Ismatec Microflex Compact Single-Channel Pump (Avantor Fluid Handling)

### Solutions

#### LB medium (1 L)

20 g Difco LB Broth Lennox (BD, REF: 240230)

Dissolve in 1 L MilliQ or ddH_2_O

Autoclave

Store at RT

**Note:** In terms of LB medium selected for this protein expression protocol, the quality of the LB broth used seems to be critical for achieving high yields of recombinant protein. We highly recommend using high quality LB broth (listed above, or your lab’s trusted ‘go-to’ LB).

#### 5 M Imidazole, pH 8.0 (100 mL)

34.04 g Imidazole, 99% (Thermo Scientific, Cat#: A10221.36)

Dissolve in 70 ml of MilliQ or ddH_2_O

Adjust pH to 8.0 with HCl

Bring volume to 100 ml with MilliQ or ddH_2_O

Store at 4°C for up to 1 month

#### IPTG stock solution, 1 M (10 mL)

2.383 g IPTG (Isopropyl-β-D-thiogalactopyranoside), Dioxane Free (RPI, Cat#: I56000)

Dissolve in 70 ml of MilliQ or ddH_2_O

Bring volume to 10 ml with MilliQ or ddH_2_O

Filter using a 0.22-µm filter membrane

Store at -20°C for up to 1 year

**Note:** Ensuring the quality of an activity of the IPTG used is also a key factor in determining protein yields for this purification protocol. We suggest freezing aliquots in ‘single use’ sizes.

#### Lysis buffer (200 mL)

10 ml Tris (1 M), pH 8.0 (Invitrogen, REF: AM9856)

40 ml 5 M NaCl (Invitrogen, REF: AM9759)

1.2 ml 5 M Imidazole, pH 8.0 (see recipe)

0.6 g Tween 20 (Sigma-Aldrich, Cat#: P9416-100ML)

69.9 µl 2-Mercaptoethanol (Sigma-Aldrich, Cat#: 63689-100ML-F)

Fill to final volume with MilliQ or ddH_2_O

Filter using a 0.22-µm filter membrane

Store at 4°C

*Final: 50 mM Tris, pH 8*.*0, 1 M NaCl, 30 mM Imidazole, 0*.*3% (w/v) Tween 20, 5 mM 2-Mercaptoethanol*

#### (optional) Endotoxin removal wash buffer 1 (400 mL)

20 ml Tris (1 M), pH 8.0 (Invitrogen, REF: AM9856)

24 ml 5 M NaCl (Invitrogen, REF: AM9759)

2.4 ml 5 M Imidazole, pH 8.0 (see recipe)

0.4 g Triton X-114 (Sigma-Aldrich, Cat#: X114-100ML)

Fill to final volume with MilliQ or ddH_2_O

Store at 4°C

*Final: 50 mM Tris-HCl, 300 mM NaCl, 30 mM Imidazole, 0*.*1% (w/v) Triton X-114, pH 8*.*0*

#### (optional) Endotoxin removal wash buffer 2 (200 mL)

10 ml Tris (1 M), pH 8.0 (Invitrogen, REF: AM9856)

12 ml 5 M NaCl (Invitrogen, REF: AM9759)

1.2 ml 5 M Imidazole, pH 8.0 (see recipe)

Fill to final volume with MilliQ or ddH_2_O

Store at 4°C

*Final: 50 mM Tris-HCl, 300 mM NaCl, 30 mM Imidazole, pH 8*.*0*

**Note:** As mentioned below, endotoxin removal is advised when RNAs treated with this RNase R will be used either in cell culture experiments and is likely required if the RNA is intended for animal experiments.

#### Wash buffer (250 mL)

12.5 ml Tris (1 M), pH 8.0 (Invitrogen, REF: AM9856)

15 ml 5 M NaCl (Invitrogen, REF: AM9759)

1.5 ml 5 M Imidazole, pH 8.0 (see recipe)

87.4 µl 2-Mercaptoethanol (Sigma-Aldrich, Cat#: 63689-100ML-F)

Fill to final volume with MilliQ or ddH_2_O

Filter using a 0.22-µm filter membrane

Store at 4°C

*Final: 50 mM Tris, pH 8*.*0, 300 mM NaCl, 30 mM Imidazole, 5 mM 2-Mercaptoethanol*

**Note:** Adding 2-Mercaptoethanol as a reducing reagent to both the lysis and wash buffers is critical to minimize nonspecific protein binding to RNase R during protein binding and washing steps. However, we highly recommend that both the elution and dialysis buffers have no reducing reagents in order to measure protein concentration directly using the spectrophotometer.

#### Elution buffer (100 mL)

5 ml Tris (1 M), pH 8.0 (Invitrogen, REF: AM9856)

6 ml 5 M NaCl (Invitrogen, REF: AM9759)

10 ml 5 M Imidazole, pH 8.0 (see recipe)

Fill to final volume with MilliQ or ddH_2_O

Filter using a 0.22-µm filter membrane

Store at 4°C

*Final: 50 mM Tris, pH 8*.*0, 300 mM NaCl, 500 mM Imidazole*

#### Dialysis buffer (1000 mL)

50 ml Tris (1 M), pH 8.0 (Invitrogen, REF: AM9856)

60 ml 5 M NaCl (Invitrogen, REF: AM9759)

630 g Glycerol, ACS Grade, (RPI, Cat#: G22025)

Fill to final volume with MilliQ or ddH_2_O

Store at 4°C

*Final: 50 mM Tris, pH 8*.*0, 300 mM NaCl, 50% (v/v) Glycerol*

#### 10x RNase R reaction buffer (10 mL)

2 ml 1 M Tris-HCl, pH 7.5 (Invitrogen, Cat#: 15567-027)

5 ml 2 M KCl (Invitrogen, Cat#: AM9640G)

25 µl 1 M MgCl2 (Invitrogen, Cat#: AM9530G)

Fill to final volume with MilliQ or ddH_2_O

Store at -20°C

Final: 200 mM Tris-HCl (pH 7.5), 1 M KCl, 2.5 mM MgCl_2_

#### Before you begin

##### Validate the sequences of the expression construct by sequencing

1. We strongly advise validating the sequences of <u>all plasmid stocks</u> regardless of whether they are received from reputable external sources or developed in house.
  a. We also suggest using whole plasmid sequencing as opposed to sanger sequencing as a complete sequence can identify issues with the plasmid that would be missed by a targeted approach.
    i. Numerous vendors offer whole plasmid sequencing for affordable rates.
2. Ensure that you have your materials gathered and have all the required reagents and buffers on hand.
  a. Selected solutions can be prepared beforehand and stored as indicated above. These include:
    i. LB media
    ii. 5M Imidazole
    iii. 1M IPTG stock solution
    iv. 100 mg/ml Ampicillin (we purchase this pre-made)
    v. RNase R Reaction Buffer
  b. Other solutions should be made fresh either on the day of (or one day before) your preparation. These include:
    i. Lysis Buffer
    ii. Wash Buffer
    iii. Elution Buffer
    iv. Dialysis Buffer
    v. (optional) Endotoxin removal wash buffer 1
    vi. (optional) Endotoxin removal wash buffer 2

#### Day 1

##### Transformation of chemically competent E. coli cells

3. Thaw chemically competent *E. coli* cells on ice.
  a. We regularly use –and this protocol is optimized for– the One Shot BL21 Star (DE3) strain from Invitrogen but others can be used per lab preferences.
4. Transform the *E. coli* according to a protocol appropriate for the strain.
  a. Add 50 ng of plasmid DNA encoding *E. coli* Rnase R to 10 µl of chemically competent *E. coli* cells.
    i. Our plasmid for WT truncated RNase R (His6-V5-RNaseR-87-725) is Addgene ID #254213 and the protocol is optimized to purify this protein.
      1. **Note:** The reaction buffer we describe is also optimized for this protein.
    ii. His6-V5-RNaseR-87-725-D280N (87-725-D280N mutant) Addgene ID #254212
    iii. His6-V5-RNase R (Full Length) Addgene ID #254214
    iv. His6-V5-RNaseR-D280N (Full Length-D280N mutant) Addgene ID #254215
    v. His6-V5-RNaseR-87-725-D278N (87-725-D278N mutant) Addgene ID #254216
    vi. His6-V5-RNaseR-87-725-D280A (87-725-D280A mutant) Addgene ID #254217
    vii. His6-V5-RNaseR-87-725-D278A (87-725-D278A mutant) Addgene ID #254218
  b. Put it on ice for 5 min.
  c. Heat shock for 45 s at 42°C using a pre-warmed MultiTherm Shaker.
    i. Shaking is not needed for this step.
    ii. **OPTIONAL:** add some water into the well to maximize heat shock efficiency.
  d. Add 100 µl of SOC Outgrowth Medium (without antibiotic) and gently pipette the cells
    i. **OPTIONAL:** Add 500 µl of SOC Outgrowth Medium (without antibiotic) and gently pipette the cells.
      1. Allow the cells to recover in selection-free media @ 37°C for 30 minutes
      2. This step is **Optional** for the constructs listed above and others plated onto AMP plates but is needed if the plasmid has a Kanamycin resistance marker.
  e. Spread cells on an LB agar plate with 100 µg/ml Ampicillin (or alternate antibiotic matching the resistance gene on the plasmid).
    i. We recommend using autoclaved glass beads for this, but traditional spreaders are also an option.
  f. Incubate the plate overnight at 37°C in a bacterial incubator.
5. Remove the plates from the incubator and ensure that single colonies are visible. Store at 4°C until needed.
  a. If plate seams are wrapped properly with parafilm, then these plates can be stored at 4°C for up to 1 month and be used for multiple preparations.
  b. When the plates are no longer needed, make sure to inactivate the *E. coli* and dispose of them using the procedures established by your institution for biohazardous waste.

#### Day2

##### Growing a ‘starter culture’ of the transformed E. coli

6. Select an isolated colony from the plate and inoculate into 100 ml LB medium with 50 µg/ml Ampicillin in a 250 ml Erlenmeyer flask.
  a. Ideally a single colony will be used, but multiple colonies are also OK.
7. Clamp the flask into an appropriate holder in a (pre-warmed to 37°C) shaking incubator.
8. Grow overnight at 37°C with shaking at 220 rpm.

#### Day3

##### Protein expression

9. Transfer 10 ml of the overnight starter culture into 1 L of LB with 50 µg/ml Ampicillin (pre-warmed to 37°C in a 2.5L culture flask)
  a. We prefer using TUNAIR Polypropylene Erlenmeyer Flasks for this step, but other equivalents are also acceptable.
  b. The remainder of the starter culture can be stored at 4°C for several days.
    i. After the procedure is completed, ensure that you inactivate and dispose of this culture using procedures established by your institution for this type of biohazardous waste.
10. Grow at 37°C with shaking at 200 rpm until OD_600_ reaches a value between 0.3 to 0.4.
  a. *Starting at 2 hr after inoculation, remove a small sample of the culture and measure OD at 600 nm using cuvettes and a spectrophotometer*.
  b. *Depending upon the reading, repeat the OD check every 10-20 minutes*
  c. *This step typically takes 2 to 3 hr*.
11. Reduce shaker temperature to 18°C.
12. Allow the cultures to equilibrate to the new temperature by growing them for 1 hr at 18°C with shaking at 200 rpm.
13. Induce expression of protein by adding IPTG to a final concentration of 0.5 mM.
14. Incubate overnight at 18°C with shaking at 200 rpm.
  a. We usually target an induction time of 16-20 hr.

#### Day4

##### Harvest the cells

15. Add 1 drop of Antifoam 204 reagent with a disposable graduated transfer pipets and swirl the cells.
16. Transfer the culture into bottles for centrifugation and ensure that they are properly (within 0.1 g) balanced.
17. Harvest cells using a high capacity floor-model centrifuge for 10 min at 5000 x *g*, 4°C.
  a. We used the Avanti JXN-26 High speed centrifuge with the J-LITE JLA-8.1000 Fixed-Angle Aluminum Rotor.
    i. Alternatively, other types of centrifuges and/or rotors with smaller bottles can be used.
    ii. However, centrifugation conditions will need to be tailored to your system.
18. Decant the supernatant and transfer the cell pellets into a 50-ml conical tube using a silicone spatula.
  a. Make sure to inactivate and dispose of the spent media according to proper protocols established by your institution for this type of biohazardous liquid waste.
19. Use a balance tared to an empty 50 ml conical tube, check the mass of bacteria collected.
  a. Expect ∼4-5 g of *E. coli* per liter of LB medium
    i. If significantly more cells are collected, that suggests that protein has likely not been induced properly.
      1. If this occurs, the IPTG is the likeliest culprit. We advise going back to step #6 and trying again with a fresh aliquot of (or freshly-made) IPTG.
20. Freeze the cell pellets.
  a. The frozen cell pellets can be stored at -80°C for several months.
  b. Alternatively, the freezing step can be skipped and you can proceed to protein purification immediately by going to step 22 below.
    i. However, if you proceed to purification, the protocol should not be stopped until the dialysis step (#42 below) is reached.

#### Day 5

##### Lyse the cells and prepare the protein

21. Remove the cell pellets from the -80°C freezer and thaw the cells.
  a. We will place the tubes in a beaker with ∼100ml of room temperature water for this step
  b. Pellets typically thaw in ∼5-10 minutes.
22. Add cold lysis buffer to the 50-ml tube until the buffer reaches 30 ml.
  a. We typically don’t measure an exact volume or mass the tube at this step. Just adding lysis buffer until the 30 ml line on the conical tube is sufficient.
23. Resuspend the pellet with vigorous vortexing.
  a. This generally takes several minutes.
  b. You want to resuspend the cells as well as possible and break up any larger clumps; however, very small cell clusters will not significantly reduce the yield of the protocol.
24. Sonicate cells on ice while putting the 50-ml tube on ice with 35% amplitude for 2 cycles. Each cycle is 3 min of sonication -5 s on, 5 s off-with 3 min of rest between cycle.
  a. *This protocol is optimized for a 6mm horn using a Pulse 150 UltraSonic Homogenizer (Benchmark Scientific)*.
    i. *Ensure that the power setting on the sonicator is set for the proper horn size*.
  b. *Please note: Different sonicators and horns will likely require different settings. Be sure to use optimized sonication conditions for your equipment*.
25. Transfer the sonicated lysates into appropriate centrifuge tubes and ensure that they are balanced to within 0.1g.
  a. We use 50 mL screw-capped (commonly called ‘Oakridge’) tubes listed in materials above
26. Centrifuge the cell lysate using a high speed centrifuge for 45 min at 30,000 x *g*, 4°C.
  a. We used the Avanti JXN-26 High speed centrifuge with JA-25.50 Fixed-Angle Rotor
27. Transfer cleared lysate into a clean 50-ml conical tube.
  a. Inactivate and dispose of the insoluble pellet using procedures established by your institution for biohazardous waste.

#### Purify the protein via FPLC

28. Equilibrate a HisTrap HP 5 mL column with 30 ml lysis buffer (flow rate 3 mL/minute) using ÄKTA Start system with added fractionator attachment.
  a. **Figure 4:** We have included a printout of the program that we use to purify the protein with this system using the Unicorn v1.3 software.
    i. The program is built using the Unicorn start v1.3 software (Cytiva)
      1. other FPLC systems will need their own version of the program with similar parameters
  b. The ÄKTA start system can be set up and used at room temperature while keeping the tubes on ice.
  c. Alternatively, the ÄKTA start can be set up inside a cold room or chromatography refrigerator.
29. Once the column is equilibrated with lysis buffer, remove the input tubing from the lysis buffer and place it into the tube with the cleared lysate.
  a. Ensure that no bubbles enter the tubing when you are moving it between tubes.
  b. Use the ÄKTA start to load the cleared lysate into the HisTrap HP 5 mL column (flow rate 3 mL/minute).
  c. Samples of the flowthrough should be collected to estimate binding efficiency.
    i. We suggest sampling at minute 3 (∼9-10 ml into sample) and minute 7 (∼20 ml into sample).
    ii. The rest of the flowthrough can be discarded.
30. **OPTIONAL endotoxin removal step**: If the RNase R yielded by this protocol is intended for purifying circRNAs that will be transfected into cells, then removing trace amounts of endotoxin from the *E. coli* is strongly recommended.
  a. Disconnect the HisTrap HP 5 mL column after protein binding step.
    i. Ensure that no bubbles enter the tubing of the ÄKTA start system or the column itself.
  b. Connect the column to a suitable pump system with clean tubing. We used a Masterflex Ismatec Microflex Compact Single-Channel Pump (Avantor Fluid Handling).
    i. Ensure that no bubbles enter the tubing or the column itself.
  c. Wash the protein-bound beads at 4°C using 300-350 mL of pre-cooled 4°C endotoxin wash buffer 1 (50 mM Tris-HCl, 300 mM NaCl, 30 mM Imidazole, 0.1% (w/v) Triton X-114, pH 8.0) to remove contaminating endotoxin.
    i. **NOTE:** This Triton X-114 removal step only works when all steps take place at 4°C.
    ii. The flow rate should be 5 mL/minute.
    iii. **NOTE:** As this step takes 60-70 minutes, it’s a good place for a short break.
  d. Wash the protein-bound beads at 4°C using 100-150 mL of pre-cooled endotoxin wash buffer 2 (50 mM Tris-HCl, 300 mM NaCl, 30 mM Imidazole, pH 8.0) to remove contaminating endotoxin.
    i. The flow rate should be 5 mL/minute.
  e. Reconnect the HisTrap HP 5 mL column to the ÄKTA start system.
    i. Ensure that no bubbles enter the column or the tubing of the ÄKTA start system.
31. Run the HisTrap HP 5 mL column using the following gradients: Wash out unbound protein with wash buffer for 16 column volumes (CV) (80 ml)
  a. NOTE: If you did the optional endotoxin removal described in step #30 above, then reducing this additional was to one column volume (5 ml) is sufficient.
  b. The flow rate should be 5 mL/minute
  c. If you are running your system at room temperature, then make sure to keep the wash buffer cool on ice.
  d. This wash fraction can go immediately to the disposal bottle
    i. Alternately, some samples can be saved to validate efficient loading of the protein onto the column and to estimate the amount of protein lost.
    ii. If wash fractions are being saved for analysis, then we suggest saving samples at early (∼2 min), middle (∼8 min), and late (∼15 min) points during the wash cycle.
32. After the wash, elute with linear gradient to 70% elution buffer in 7 CV and then step up to 100% elution buffer for 5 CV.
  a. The flow rate should be 2 mL/minute for this step.
  b. If you are running your system at room temperature, then make sure to keep the elution buffer cool on ice.
  c. Ensure that you are capturing these fractions for analysis
  d. We generally take 1 ml fractions during elution.
  e. The ÄKTA start system has a built-in monitor that can track the absorbance of the protein samples as they are eluted. Observing the trace of the eluted fractions will give you an estimate of which fractions contain the bulk of the eluted proteins.
  f. Make sure to clean the ÄKTA start system and rinse out the tubing with 6 CV of MilliQ water with 0.01% sodium azide to inhibit bacteria growth while the machine in storage mode between purification runs.
33. Prepare an electrophoresis sample from the eluted fractions to evaluate the purified protein by SDS PAGE.
  a. Mix 5 µl of samples and 25 µl of 2x Laemmli Sample Buffer.
  b. Heat the samples at 95°C for ∼5 minutes.
34. Run on a Stain-free SDS-PAGE gel (load 15 µl per lane)
  a. We prefer using the 4-20% pre-cast Stain-free gradient gels listed above so we can see proteins of all sizes to assess the purity of our purified proteins.
  b. Alternately, standard (in-house made) SDS-PAGE gels are sufficient. We suggest a 10-12% gel to visualize possible contaminants or degradation products that could occur during protein expression or purification.
35. Detect protein bands using ChemiDoc Imaging System on Stain Free Gel mode.
  a. If you are not using stain-free gels, then the gels must be stained to visualize the proteins. As the protocol produces large quantities of RNase R, a standard Coomassie-type stain (such as Coomassie Brilliant Blue) is sufficient.
    i. As an easier to use alternative to traditional Coomassie stain, we suggest using this Colloidal Blue Staining Kit (LC6025 from Invitrogen). Although initially more expensive, the ‘used stain’ can be saved and re-used for months making it more cost efficient.
36. Examine the stained gel and select the fractions that have the ∼70 kDa band corresponding to His6-V5-RNaseR-87-725.
  a. Ensure that few/no degradation products or contaminating proteins are visible below the protein band.
  b. As mentioned above, we observed that select nuclease dead mutants of RNase R were usually contaminated with residual RNA (presumably from the lysed *E. coli*).
    i. These were not visible via SDS PAGE gel.
    ii. Nucleic acid contamination can be detected by low A260/280 ratios.
37. Combine the selected fractions of RNase R for equilibration in a 15-ml conical tube.
38. Wash a dialysis membrane with MilliQ water and seal the bottom of the membrane with a closure.
39. Transfer eluted proteins into the membrane and seal the membrane with the closure.
40. Float the membrane with a floater in the pre-cold dialysis buffer.
41. Set 5-10 ml of the dialysis buffer aside in a 15 ml conical tube and store at 4°C.
42. Dialyze the proteins in 1L of Dialysis buffer overnight at 4°C with gentle stirring on a magnetic stir plate.
  a. We advise that the dialysis step last 16-20 hours.

**Figure 4:**
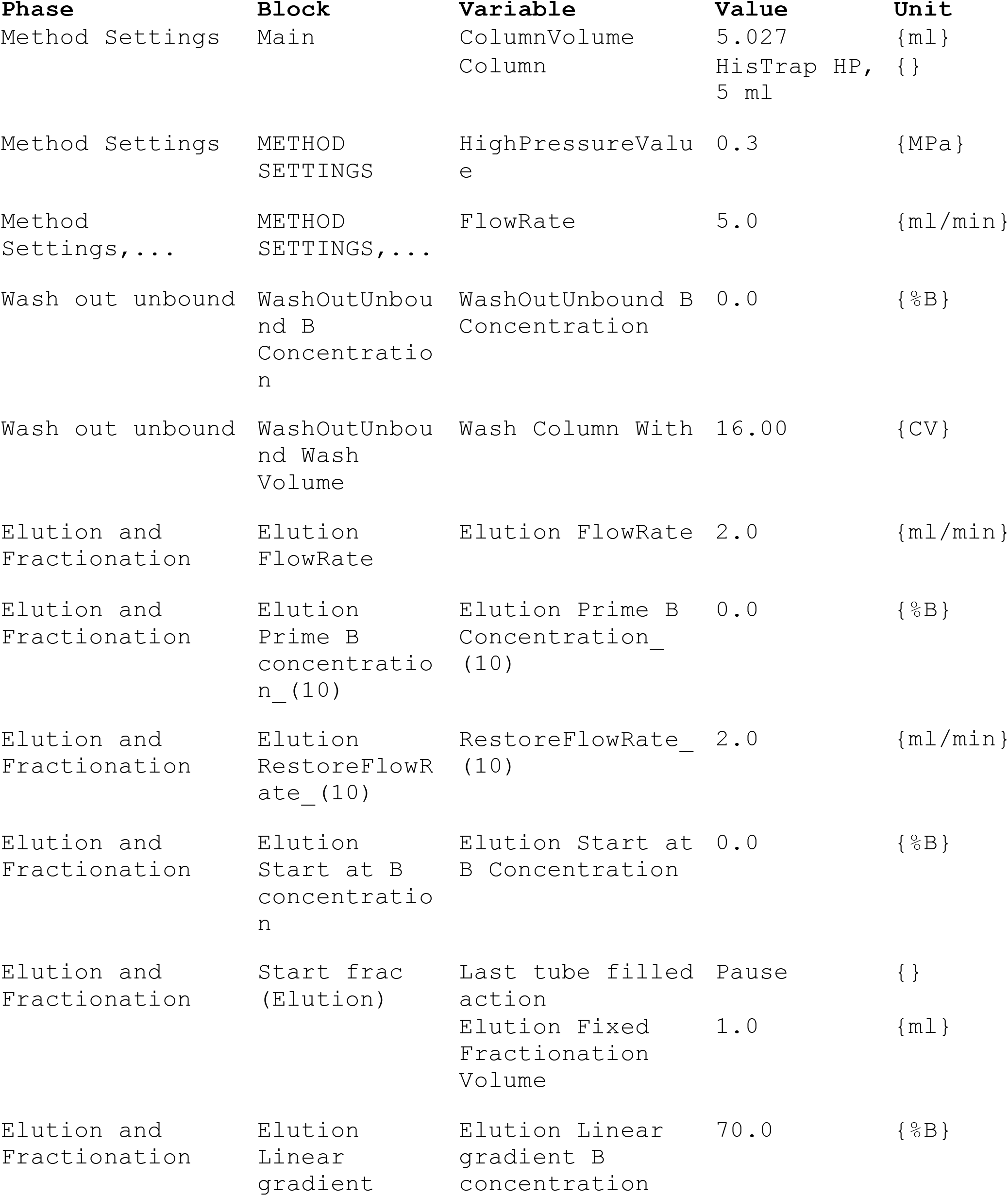
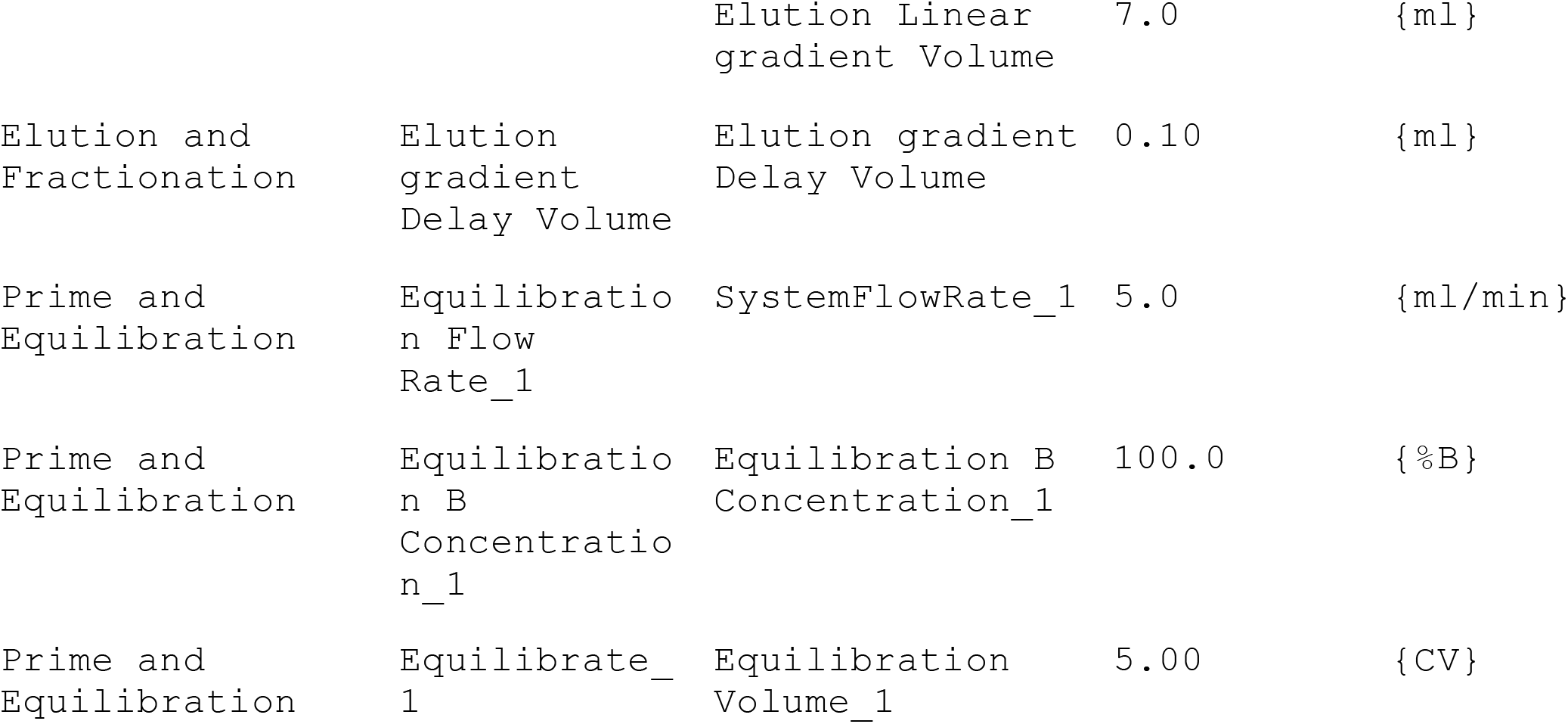
A printout of the purification program for the Unicorn v1.3 software used to purify the protein. The program here is written to be implemented on an ÄKTA start FPLC system and Frac30 fraction collector (Cytiva) running UNICORN start v1.3 software (Cytiva)

#### Day5

##### Purify the protein (continued from previous day)

43. Carefully (Remember, the dialysis buffer contains 50% glycerol, so the tubing will be slippery) remove dialysis tubing and clamps from the beaker.
44. Carefully open one side of the dialysis clips and transfer the contents of the tubing into a fresh 15 ml conical tube & place on ice.
  a. Note: the volume of the sample usually decreases by at least 50%.
45. Measure protein concentration with the spectrophotometer.
  a. For this version of RNase R (87-725) the extinction coefficient is 63,260/M/cm. That value was used to determine protein concentration by correcting protein-specific absorbance characteristics.
  b. Use some of the reserved dialysis buffer to dilute the purified RNase R to a concentration in line with your uses.
    i. Our suggested working amount is to use 0.6 µg purified RNase R per 40 µL digestion reaction. We find this amount to be functionally equivalent to one unit of (most brands of) commercially-sourced RNase R.
46. Aliquot the purified RNase R protein in convenient portions and store the protein at -20°C.
  a. With proper storage, we have documented no measurable loss of exonuclease activity for at least 14 months after purification.
47. **Please Note:** the RNase R produced using this method will retain some levels (between traces to a considerable amount) of endotoxin.
  a. If the RNase R is intended for use in reactions to ‘clean up’ circRNAs that will be transfected into cells (or encapsulated into lipid nanoparticles and delivered to animals), then the endotoxin will likely need to be removed (via the Optional step #30) to ensure that traces of endotoxin don’t induce immune responses.

### Basic Protocol 2

#### Detection of exoribonuclease activity and specificity via *in vitro* RNA decay assay

This protocol details detection of RNA degradation by recombinant RNase R using either a 5’-end FAM-labeled RNA oligonucleotide and/or a mixture of linear mRNA and circRNA in an *in vitro* RNA degradation assay. We used this assay to validate the activity and specificity of the RNase R purified using this protocol.

#### Materials

10x RNase R Reaction Buffer (200 mM Tris-HCl (pH 7.5), 1 M NaCl, 2.5 mM MgCl2, 10 mM DTT) (also see recipe above)

In vitro transcribed mRNA test sample(s) (optional) circRNA test sample(s)

(optional) 5’ end FAM-labeled RNA oligonucleotide (various suppliers)

(optional) SYBR™ Gold Nucleic Acid Gel Stain (S11494, Thermofisher)

Purified RNase R (from above)

Reference RNase R (either a commercial sample or an earlier preparation)

RNA FlashGel (Lonza or equivalent) or Urea PAGE gel for oligonucleotide substrate

**NOTE:** In our hands, the RNase R purified using this protocol <u>was highly sensitive to reaction conditions</u>. In fact, our tests showed that RNase R purified using this protocol was inactive in several commercially-provided RNase R buffers. As such, we strongly recommend that you use optimized reaction buffer (above) for eliminating linear RNAs using RNase R purified with this protocol. Although our sampling of commercially-sourced RNase R buffers was not exhaustive, our experience shows that using reaction buffers sourced from commercial RNase R will likely negatively affect the efficiency of RNA degradation.

1. Prepare a 10x stock of RNase R and 10x RNase R reaction buffer for the test.
  a. Calculate the amount of RNase R needed to make a 1mg/mL stock.
    i. Use 1x reaction buffer to make an appropriately-sized aliquot.
    ii. The target final working concentration for RNase R is 0.6 µg per 40 µL digestion reaction.
      1. This works out to a final concentration of 0.015 mg/ml RNase R in each reaction to digest 2 µg of target RNA.
    iii. This target concentration may vary slightly from batch to batch, but has remained (more or less) consistent across batches.
      1. Across several batches of RNase R purified using this protocol, on average, 0.3 µg recombinant RNase R is comparable to 1 Unit of commercially purchased RNase R.
2. Prepare the test reaction by mixing your RNA substrate(s), RNase-free water, and 10x RNase R Reaction Buffer in a tube.
  a. Make sure to use an amount that is reasonable for your detection limit. For example, this range works well with FlashGel but would be too low for a traditional agarose gel.
  b. For long ‘target RNAs’ (such as mRNAs, circRNA, etc) substrate RNA can range in concentration from 50 ng/µl up to 500 ng/µl.
    i. The amount of RNA and incubation times can be changed to suit needs.
      1. For example, we’ve validated complete digestion of 2 µg of RNA after a 30 minute digestion with 0.6 µg of purified RNase R in a 40 µl reaction volume.
  c. For labeled RNA oligos, we suggest 100 nM of RNA oligo and 100 nM of RNase R for a digestion time course from 0 to 30 minutes.
3. Add the RNase R.
4. Digest the samples for between 15 and 60 min at 37°C.
5. Stop the reaction and harvest the RNA using your preferred method.
  a. We suggest using RNA Clean and Concentrator 5 columns (Zymo)
6. Load the purified test reactions on an RNA Flashgel.
  a. You can use either the denaturing or native RNA loading buffers, but we always had best results with the formamide buffer.
  b. Alternatively, if a FAM-labeled oligo is used, then a urea PAGE gel and a ChemiDoc MP can be used to visualize the results.
  c. If using a gel without an embedded dye, then the nucleic acids can be visualized using SYBR™ Gold Nucleic Acid Gel Stain or an equivalent reagent.
7. Visualize and record the results.

### Additional Protocol 1

#### Detection of RNA binding activity via RNA electromobility shift assay (EMSA)

The EMSAs were performed according to established procedures using the 5’ FAM-labeled RNA oligonucleotide shown in **Figure 2**. The protocol is based on similar EMSA assays using DNA oligonucleotides, but was optimized for RNA and ^32^P-labeled DNA oligos were replaced with FAM-labeled RNA [7]. Briefly, purified RNase R proteins and RNA oligonucleotides were diluted to appropriate concentrations in 1x RNase R reaction buffer (see recipe above in **Basic Protocol 2**). RNA oligo and purified proteins were mixed in the indicated molar ratios ranging from equal concentrations up to 10-fold excess of protein. Samples were incubated at 37°C for 15 min. After the incubation, non-denaturing loading buffer (0.05x TBE buffer, 2.5% (v/v) Glycerol) was added to the samples and they were loaded onto pre-run 6% non-denaturing SDS PAGE gels in 0.5x TBE buffer for 30 min at 80 volts (constant voltage setting). Samples were visualized using a ChemiDoc as above.

#### Time Considerations

Growth of *E. coli* from transformation to protein expression takes 3 days. Lysis and purification of protein takes ∼4 hr. PAGE, imaging and dialysis process take ∼2 hr of hands-on time. The different activity and RNA binding assays typically take 2-4 hr.

## Acknowledgements

This work was funded as part of a larger project supported by a grant from CEPI (the Coalition for Epidemic Preparedness Innovations) where John P. Cooke (ORCID 0000-0003-0033-9138) was the Principal Investigator and DLK was the project lead for RNA development.

## Author Contributions

**WH –** Conceptualization, Methodology, Investigation, Formal analysis, Validation, Writing - Original Draft, Visualization, Resources

**DLK –** Conceptualization, Writing - Original Draft, Writing - Review & Editing, Resources, Supervision, Project administration

